# Non-apoptotic death of the *C. elegans* linker cell is primed by MYRF-1 activation of *pqn-41*/polyQ

**DOI:** 10.64898/2025.12.08.693091

**Authors:** Olya Yarychkivska, Simin Liu, Lauren Bayer Horowitz, Shiloh Newland, Peipei Wu, Saiya Mittal, Shogo Tamura, Tetiana Novosolova, David Faulkner Ritter, Yun Lu, Sevinç Ercan, Christopher Hammell, Shai Shaham

## Abstract

Linker cell-type death (LCD) is a morphologically conserved non-apoptotic cell-death process with features resembling polyglutamine-dependent neurodegeneration. In *C. elegans* development, LCD eliminates the male-specific linker cell following its long-range migration. Using single-cell mRNA sequencing of migrating and dying linker cells, we identify *myrf-1*, encoding a membrane-bound transcription factor implicated in human developmental disorders, as a key LCD regulator. MYRF-1 translocates to the linker cell nucleus during early migration and, surprisingly, its auxin-inducible degradation then, but not later, blocks LCD. MYRF-1 directly binds known LCD genes, including *pqn-41*, encoding an aggregation-prone polyglutamine protein. Deleting a *bona fide* MYRF-1-binding site within *pqn-41* promotes linker cell survival. Our findings reveal that linker cell death is primed well before cell demise takes place, temporally uncoupling death commitment and execution.

---

Ultrastructural changes accompanying **l**inker **<u>c</u>**ell-type **<u>d</u>**eath (LCD), including nuclear crenellation, loss of peripheral heterochromatin, and organelle swelling, are distinct from typical apoptotic features (*1*). Nonetheless, cell death associated with these changes is prevalent in development and in disease (*2*). In vertebrates, for example, LCD features are evident during the normal dismantling of the Müllerian duct in developing males, in cells of the developing palate, and in dying spinal cord motoneurons (*3*). A similar morphological signature is associated with dying cells in polyglutamine disease patients and in mice that model these disorders. Indeed, crenellation of striatal neuron nuclear membranes is a hallmark of Huntington’s disease (*3*) and similar nuclear envelope invaginations are widespread in autopsy samples from dentatorubral-pallidoluysian atrophy (DRPLA) patients (*4*).

While molecular players governing vertebrate LCD remain to be discovered, a genetic framework for this process has been outlined in the nematode *C. elegans* (*2*). Here, the male-specific linker cell dies by LCD at the larva-to-adult transition, following its long-range migration (*1*). Death follows a well-orchestrated sequence of events, starting with nuclear crenellation, and followed by asymmetric cell splitting, engulfment of both linker cell fragments, reduction in linker cell cytoplasmic volume, and eventual degradation (*1*, *5*) (Fig. 1A). Linker cell migration guides gonadal elongation, and linker cell death facilitates gonad-cloaca fusion, creating an open reproductive system that ensures male fertility (*6*). LCD in *C. elegans* takes about eight hours to complete, an order of magnitude longer than apoptotic events in this animal (*5*).

**Fig. 1.**
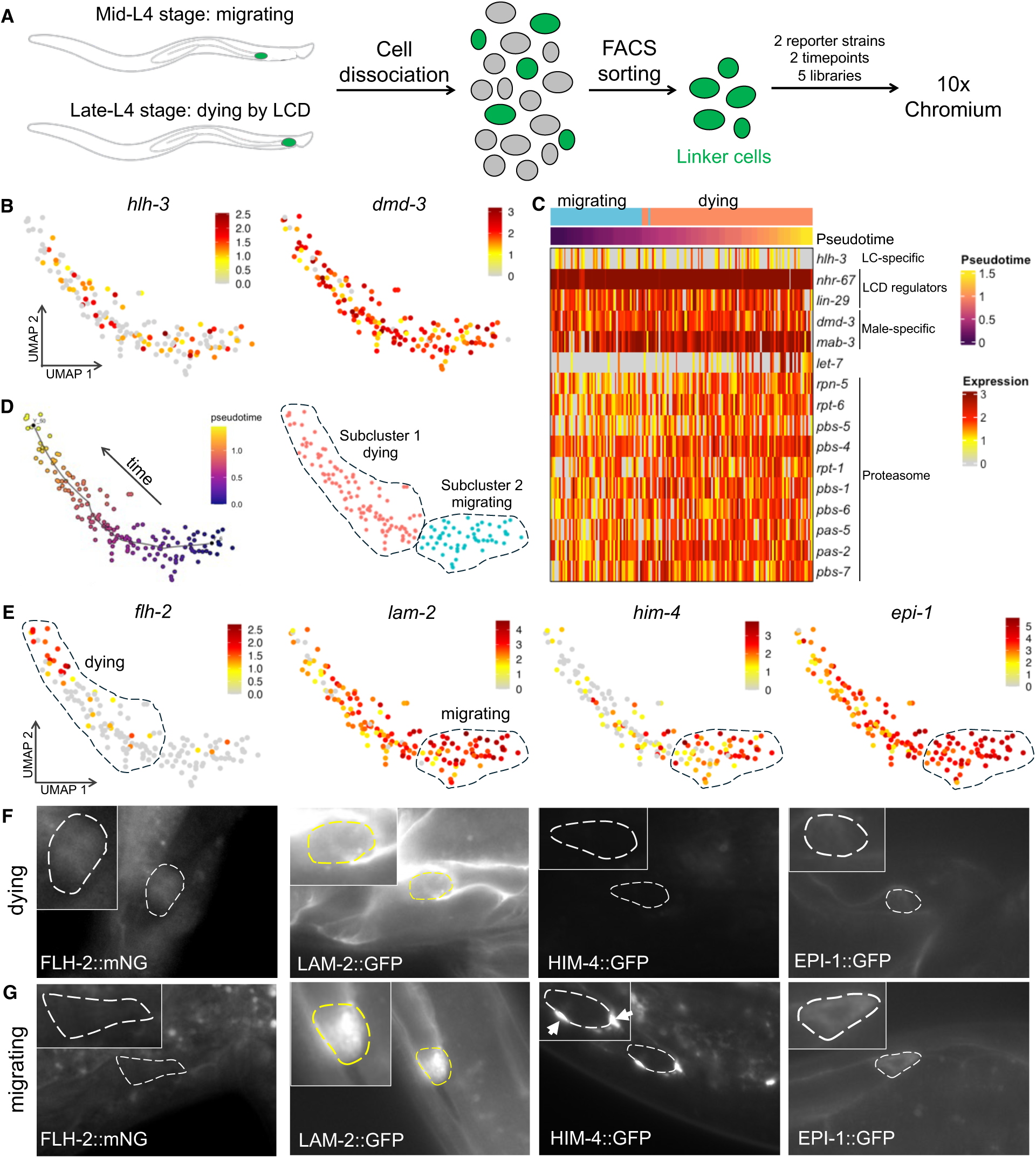
Isolation and transcriptional profiling of migrating and dying linker cells. (**A**) Schematic of linker cell isolation for 10x Chromium. (**B**) Linker cell cluster expresses LCD and male-specific genes. (**C**) Pseudotime and Monocle3 analysis identified two subclusters. (**D-F**) Expression data and *in vivo* reporter validation of corresponding genes upregulated in dying or migrating cells.

Linker cell death is molecularly distinct from apoptosis. The main apoptotic caspase, CED-3, and its regulators, are not required for linker cell death initiation (*1*), although auxiliary roles in cell dismantling and engulfment have been described (*7*). Instead, two opposing Wnt signaling pathways, developmental timing regulators, and a SEK-1/MAPKK cascade converge to control activation of a non-canonical death-promoting function of the highly conserved heat-shock responsive transcription factor HSF-1. HSF-1 activates expression of genes distinct from those induced during the heat-shock response, including *let-70*, encoding a conserved UBE2D2-like E2 ubiquitin ligase, and other components of the ubiquitin proteasome system (UPS) (*1*, *8–10*). The *C. elegans* linker cell thus provides a powerful system to study mechanisms governing LCD.

Despite progress in identifying LCD components, delineating the full complement of LCD mediators in *C. elegans* has been challenging. Linker cell survival mutants can be difficult to propagate, as they exhibit reduced fertility. Furthermore, the cell is deeply embedded within the animal (*6*) and is connected by robust adherens junctions to adjacent gonadal tissue (*1*), making its isolation for transcriptomic studies challenging. A previous study used laser microdissection to isolate a handful of migrating linker cells and identified some expressed genes within these. Nonetheless, the resulting transcriptomes are unlikely to be comprehensive and may be contaminated by mRNAs originating in neighboring spermatids and perhaps in other somatic gonadal cells (*11*).

In this study, we report the development of a method to purify hundreds of intact migrating and dying linker cells. We use this pipeline to identify single-cell transcriptomes of a continuum of linker cell developmental stages spanning the migration-to-death transition and resolved to within 1-2 minutes. Our data suggest characteristic transcriptional changes accompanying LCD onset, involving downregulation and upregulation of many genes. We use this data to identify new LCD regulators, including the conserved membrane-bound transcription factor MYRF-1, implicated in human developmental disorders and in cancer (*12–19*). We show that in animals lacking MYRF-1, the linker cell completes its migration but fails to initiate LCD. MYRF-1 functions, surprisingly, not at the time of LCD, but much earlier, during the L2-to-L3 larval transition, when the cell makes a posterior turn in its migration path. MYRF-1 translocates from the cell membrane to the nucleus during this turning event; binds promoter regions of several LCD regulators, including *pqn-41,* a gene encoding an endogenous polyQ protein required for LCD; and controls *pqn-41* expression. Our findings demonstrate that linker cell death commitment is temporally uncoupled from LCD execution and raise the possibility that similar priming mechanisms operate in other developmental and disease contexts.

## Results

### A developmental transcriptome of the linker cell as it initiates LCD

To identify genes required for LCD, we sought to isolate individual linker cells as they undergo the migration-to-death transition and to characterize their transcriptomes. We examined two linker cell reporter strains. In one, the linker cell is marked with a *mig-24p::gfp* reporter, also expressed in the hermaphrodite distal tip cells (DTCs) (*20*). In the other, a combination of *mig-24p::gfp* and *nhr-67::mCherry* reporters marks the linker cell more specifically (fig. S1A). Both strains are also homozygous for the *him-5(e1490)* allele, which increases the incidence of males from 0.2% to ∼30% of the population (*21*). Strains were bleach synchronized (*22*) to generate L1-stage larvae that were allowed to grow for about 36 hours at 20°C to the mid-L4 larval stage. The position of the linker cell was then monitored until most linker cells were either executing the final leg of their posterior migration or reached the tail and initiated LCD (Fig. 1A). Animals at each of these two timepoints were collected and subjected to carefully monitored enzymatic and mechanical disruption (Methods). Rare linker cells, predicted to comprise only 0.03% of the total cell suspension, were isolated using fluorescence activated cell sorting (FACS) and promptly processed for single-cell RNA sequencing (scRNA-seq) using the 10x Chromium platform (Fig. 1A). To preserve temporal and strain information, samples from different timepoints and strains were processed independently.

We generated five sequencing libraries: two from migrating-stage and three from dying-stage linker cells (fig. S1B). Standard quality control measures were implemented to eliminate ambient RNA, cell doublets, and cells with high mitochondrial transcript content (Methods). Clustering analysis, UMAP plotting, and examination of genes with known expression patterns allowed us to unambiguously identify 155 linker cells (Fig. 1B), 2,230 DTCs, as well as additional cell types (fig. S1B). Supporting our assignment, cells within the linker cell cluster express the lineage-specific transcription factor gene *hlh-3* (*20*); the male-specific genes *dmd-3* and *mab-3*; the LCD regulators *nhr-67* and *lin-29* (*1*, *9*); and other expected gene transcripts (Fig. 1B,C; fig. S1C,D, Data S1, S2). DTC cells were assigned to two distinct clusters based on expression of *hlh-2*, *hlh-12*, *lin-32*, *apx-1*, *lag-2*, and *ina-1* (fig. S1D) (*20*). The linker cell-to-DTC ratio in our final analysis sample (1:14.4) is considerably lower than predicted based on male abundance in the population (1:4.7). This discrepancy is likely due to difficulties in disrupting linker cell junctions to neighboring cells.

### Identification of migrating and dying linker cell populations

A combined UMAP projection of linker cells isolated from all five experiments reveals a continuous transcriptional trajectory (Fig. 1D). To infer temporal relationships among cells, we performed a pseudotime analysis, which orders cells based on transcriptional similarity (*23–26*). The resulting trajectory reflects the migration-to-death progression, as indicated by the increased expression of *let-7* miRNA (Fig. 1C), a key regulator of the larva-to-adult transition (*27*), along the pseudotime axis (see below; Data S3). Because animals collected at the migrating and dying timepoints exhibit some variability over linker cell development stages, their combined transcriptomes span a smooth distribution over the two-hour sample collection period, providing an effective temporal resolution of ∼1–2 minutes.

To further validate our temporal assignments, we applied a clustering algorithm to the combined linker cell dataset, revealing two subpopulations that appear to correspond to migrating and dying linker cells (Fig. 1D) (*23–26*). Differential gene expression (DEG) analysis (*28*) identified 149 and 153 enriched gene transcripts in each subcluster, respectively (Data S3). Consistent with our assignments, *let-7* miRNA is induced in the dying cell cluster, as are transcripts encoding proteasome components, such as *rpn-5* and others (Fig. 1C, Data S3,), consistent with our previous studies documenting increased expression of *rpn-3*, a 26S proteasome subunit, during LCD (*8*). Furthermore, we showed that an mNeonGreen reporter fusion to *flh-2*, a gene enriched in the dying cell cluster, is induced *in vivo* upon LCD initiation (Fig. 1E-G). Genes enriched in the migrating cell cluster include the known cell migration genes *him-4*, *epi-1*, and *lam-2* (*29*), and reporter fusions to these genes are expressed *in vivo* in migrating but not dying linker cells (Fig. 1E-G). Finally, to capture low abundance but biologically relevant transcripts, we developed a rare gene analysis pipeline (Methods) to identify significantly changing genes expressed in 5-20% of cells comprising the linker cell cluster. This analysis identified 173 additional genes upregulated in dying cells and 73 additional genes upregulated in migrating cells (Data S4).

We also performed gene module analysis (*23–26*) on our pseudotime-ordered cells to identify coordinated gene expression programs along the linker cell developmental trajectory. This revealed two distinct transcriptional programs: a Migration Module upregulated in migrating cells and downregulated in dying cells (fig. S2A,B; Data S3), and an LCD Module, which shows the reciprocal pattern (fig. S2A,B; Data S3).

We used the pseudotime, DEG, and gene module analyses to generate high confidence lists comprising 297 genes upregulated in dying cells and 213 genes enriched in migrating cells (Fig. 2A; fig.S2C; Data S3). The LCD list contains the known LCD regulators *let-7*, *lit-1*, *nhr-67*, and *ebax-1*, among others (*1*, *8*, *9*) (Horowitz et al., in prep; Fig. 2A; fig. S1C, Data S3). Gene Ontology (GO) analysis of this list reveals enrichment of proteasome components as well as genes associated with vesicle-mediated transport (Fig. 2A). Migrating list GO terms reflect enrichment of genes involved in translation, RNA biosynthesis, and ribosome biogenesis, as well as components associated with ribosomes, microtubule cytoskeleton, and mitochondrial membrane. Together, these data uncover new gene networks that likely direct linker cell migration and death.

**Fig. 2.**
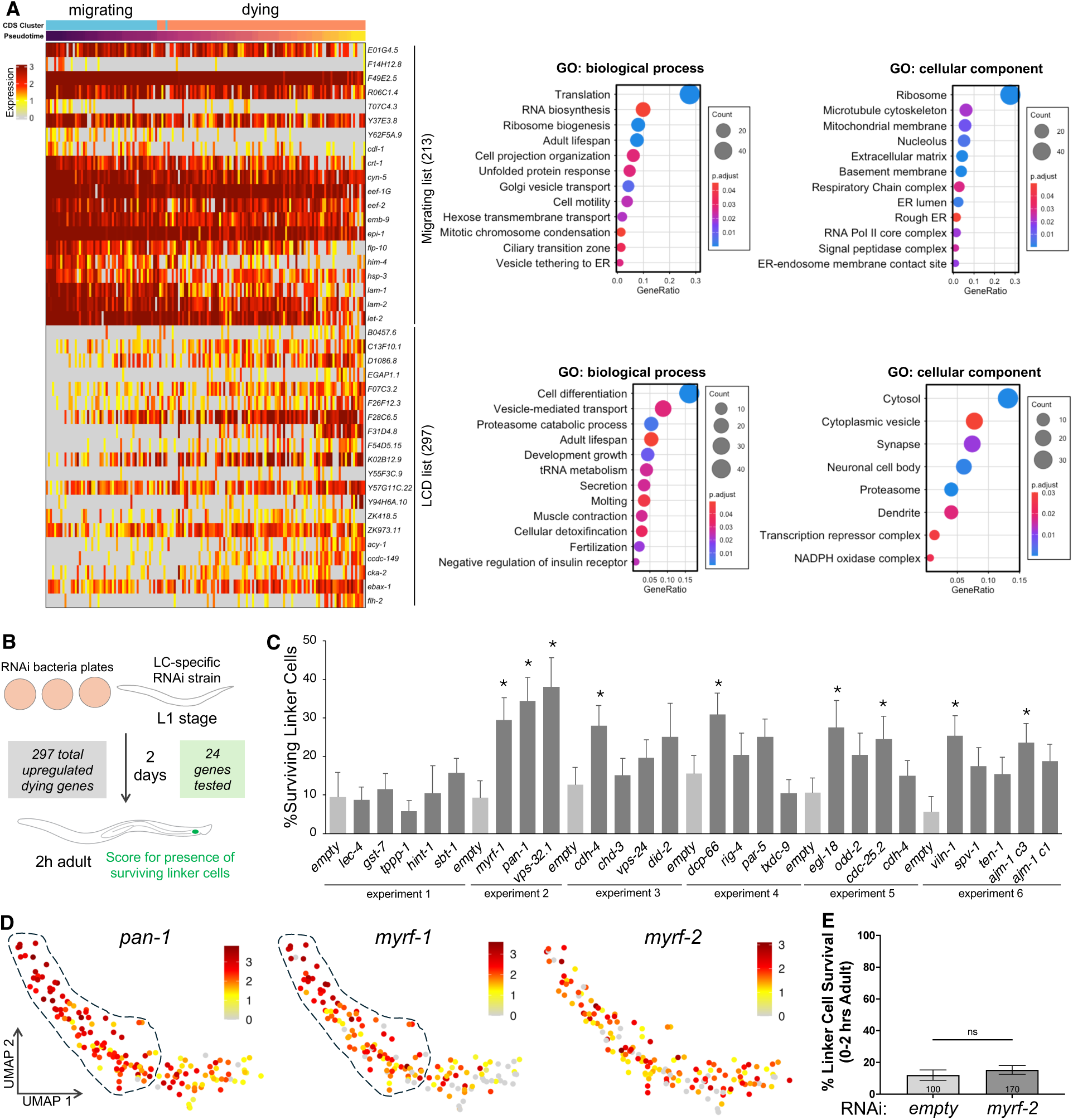
Analysis of scRNA-seq data and identification of novel LCD regulators. **(A)** Top 20 genes from Migrating and LCD lists showing gene expression changes along pseudotime (purple – start, yellow – end), and their Gene Ontology analysis. **(B)** Schematic of a LC-specific RNAi screen. **(B)** Identification of novel LCD regulators from LC-specific RNAi screen. (**C**) Screen hits expression in the linker cell cluster. (**D**) *myrf-2* RNAi has no effect on LCD.

### RNAi screen identifies previously unknown LCD genes

Genes expressed in the dying linker cell might represent previously unknown LCD regulators. We therefore used linker cell-specific RNAi, which allows assessment of genes even if their systemic loss leads to lethality, to interrogate the functions of 24 genes whose expression is either increased or unperturbed as the linker cell dies (Fig. 2B; Data S1-S4). Remarkably, nine of the RNAi clones we tested induce aberrant linker cell survival two hours after the larva-to-adult molt (Fig. 2C). This nearly doubles the known gene complement required for LCD in *C. elegans* (*2*). These new regulators include *vps-32.1*, involved in vesicle transport (*30*), *cdh-4*, encoding a cadherin-related protein involved in cell adhesion (*31*), *dcp-66*, a predicted member of the NuRD transcriptional complex (*32*), *egl-18*, encoding a GATA-like transcription factor (*33*), *cdc-25.2*, encoding a conserved cell cycle regulator (*34*), *viln-1*, encoding the *C. elegans* cytoskeletal protein villin (*35*), and *ajm-1*, encoding a major *C. elegans* adherens junction protein (*36*). These results suggest that several cellular functions, including vesicle-mediated transport, cell adhesion, and transcription, are necessary for linker cell death.

### MYRF-1 is cell-autonomously required for linker cell death

In our RNAi screen we also identified *myrf-1*, encoding a conserved membrane-tethered transcription factor (*19*) as a possible linker cell death gene (Fig. 2C). Mammalian MYRF was first identified as a regulator of oligodendrocyte differentiation (*37*). In humans, MYRF is essential for viability, and heterozygous mutations are associated with a panoply of rare developmental disorders (*12–16*, *19*) and with cancer (*17*). In *C. elegans*, MYRF-1 regulates developmental progression and loss of *myrf-1* results in early larval arrest (*38*, *39*). To validate our scRNA-seq findings showing *myrf-1* upregulation in dying cells (Fig. 2D), we examined animals in which MYRF-1 is endogenously tagged with GFP. We observed expression of the tagged protein in the linker cell and confirmed induction of expression at the late larval L4 stage, preceding the onset of LCD (fig. S2D). To further confirm the effects of *myrf-1* loss on the linker cell, we repeated the *myrf-1* RNAi experiment (n=3 repeats) and demonstrated that although linker cell migration is unaffected in knockdown animals, the linker cell, nonetheless, ectopically survives in 2 h and 24 h adults (fig. S2E).

We also developed a method to degrade MYRF-1 protein specifically in the linker cell using an auxin-inducible degron (AID) (*40*). Three strains were endogenously engineered to express either AID-GFP-MYRF-1 N-terminal fusion protein, MYRF-1 containing GFP-AID inserted after amino-acid 171, or MYRF-1 containing mKate2-AID inserted at the same location (Fig. 3A, fig. S3A) (*39*). Into these animals, we introduced a *mig-24p::TIR1* transgene, expressing the TIR1 E3 protein, which mediates degradation of AID-tagged proteins in the presence of auxin specifically in the linker cell (Fig. 3B). In the absence of auxin, animals are viable, develop normally, and linker cell death proceeds in a timely manner (Fig. 3C water control, fig. S3A). Upon continuous exposure to auxin, animals remain viable, exhibit no developmental defects, and display mostly normal linker cell migration (Fig. 3D) (∼20% of surviving linker cells have a slight migration defect, fig. S3B, C); however, the linker cell now survives inappropriately in most animals (Fig. 3C, fig. S3A). Surviving linker cells fail to compact their cytoplasm, harbor normal cytoplasmic organelles, and do not exhibit the deep nuclear crenellations characteristic of LCD (Fig. 3E), while linker cells from control (water-treated) animals display deformed nuclei and swollen ER (fig. S3D).

**Fig 3.**
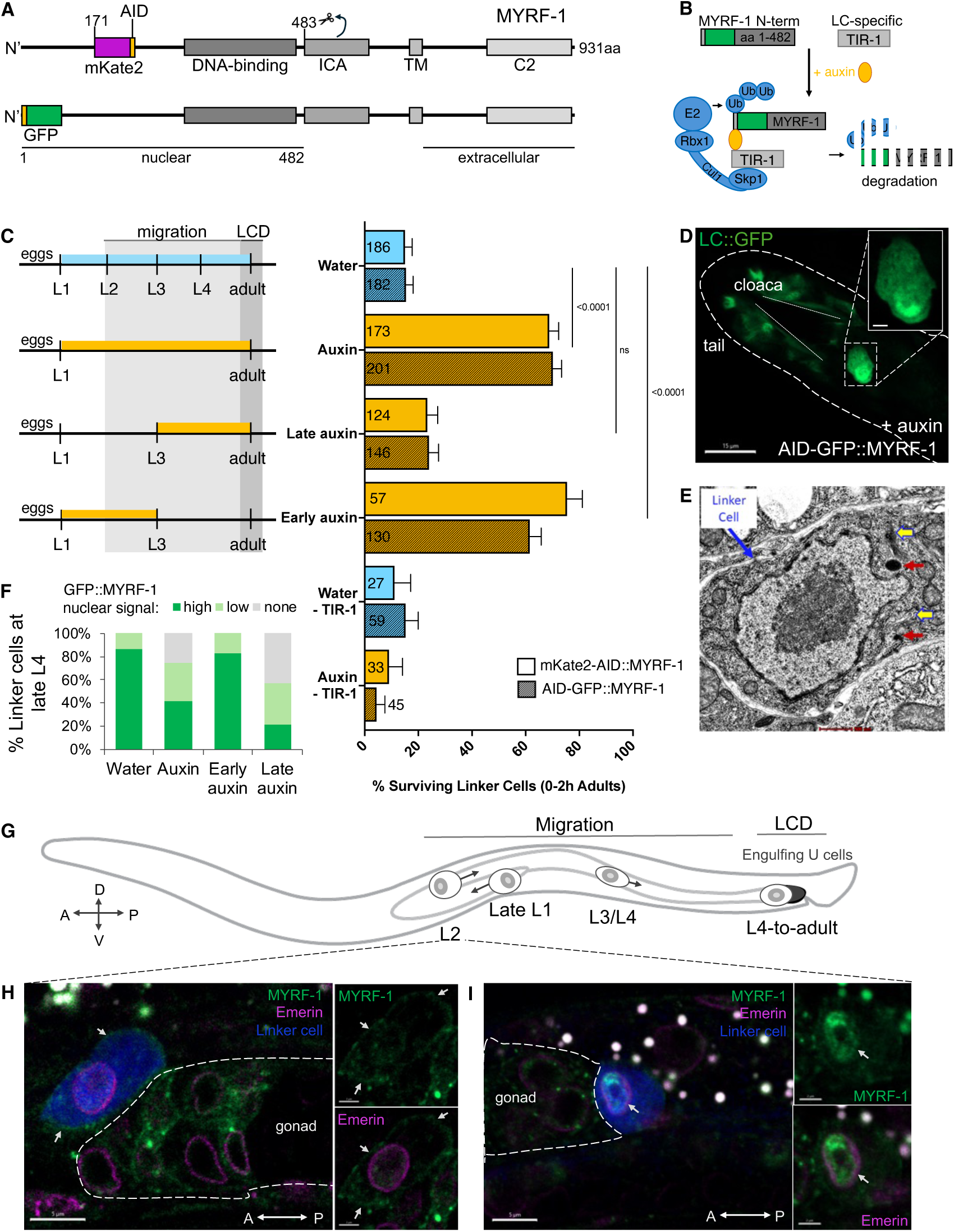
MYRF-1 functions during early linker cell migration to promote linker cell death. **(A)** MYRF-1 structures with corresponding inserted fluorescence and AID tags. **(B)** Schematic of a linker cell-specific AID experiment. **(C)** Depletion of MYRF-1 during specific developmental timeframes. Importantly, the linker cell is only born in the L2 stage, thus L1-L3 auxin treatment actually reflects L2-L3 treatment of the linker cell **(D)** Surviving linker cell in a MYRF-1 depleted young adult male. **(E**) Ultrastructural studies of a surviving linker cell in a MYRF-1 depleted adult male. **(F)** Quantification of linker cells with the nuclear MYRF-1 signal in late L4 stage in different AID conditions used in **(C). (G)** Schematic of linker cell migration during the animal larval development. **(H-J)** Confocal imaging of N-terminally GFP-tagged MYRF-1 during late L2 migration.

Finally, we found that like *myrf-1* mRNA, mRNA encoding PAN-1, a transmembrane interacting partner of MYRF-1 (*41*), is upregulated in the linker cell as it dies (Fig. 2D). Importantly, linker-cell-specific RNAi against *pan-1* also results in inappropriate linker cell survival at 2 h and 24 h adults (Fig. 2C, fig. S2E). Thus, multiple independent lines of evidence support a cell autonomous function for MYRF-1, likely working with PAN-1, in linker cell demise.

In addition to MYRF-1, the *C. elegans* genome also encodes another MYRF-related protein, MYRF-2, required for developmental maturation of DD motoneurons (*42*). Our scRNA-seq data suggest that while *myrf-2* mRNA is expressed in the linker cell, it is not upregulated in dying cells (Fig. 2D). Furthermore, linker cell death proceeds normally in *myrf-2*(RNAi) animals (Fig. 2E). Thus, MYRF-dependent processes controlling linker cell death are likely distinct from those controlling neuronal development.

### MYRF-1 primes the linker cell for death in early larval development

*myrf-1* mRNA upregulation as the linker cell begins to die raised the possibility that MYRF-1 acts during this transition. To test this, we exposed *myrf-1*::GFP-AID animals to auxin over various time intervals during development and quantified aberrant linker cell survival. Surprisingly, depleting MYRF-1 protein just prior to and during the larva-to-adult transition has no effect on linker cell death (Fig. 3C, fig. S3A). By contrast, removing MYRF-1 >15 hours earlier, at the L2-L3 larval stages, when the linker cell just completes its anterior migration and begins to move posteriorly, robustly inhibits linker cell death (Fig. 3C). This block is maintained even 24 hours later (fig. S3E), supporting the notion that MYRF-1 loss does not delay LCD progression, but is rather required for LCD onset. To exclude the possibility that early depletion of MYRF-1 affects its abundance at later stages, we examined MYRF-1 expression in L4 animals subjected to an early MYRF-1 depletion-restoration regimen. We found comparable levels of AID-GFP::MYRF-1 protein in these L4 animals and in animals untreated with auxin (Fig. 3F). These findings are consistent with the idea that MYRF-1 is required only during early development to specify linker cell death and that *myrf-1* mRNA upregulation immediately preceding the larva-to-adult transition is not relevant for LCD execution. Further supporting an early role for *myrf-1* in LCD regulation, we found that as the linker cell switches migration direction at the L2-to-L3 transition, MYRF-1 translocates from the plasma membrane to the nucleus (Fig. 3H, I).

Our studies therefore reveal that MYRF-1 primes the linker cell to die far in advance of death initiation and suggest that MYRF-1-mediated transcription at this early stage is likely required for later initiation of LCD.

### MYRF1 binds to and functionally regulates *pqn-41*/polyQ gene expression for LCD

The early action of MYRF-1 suggests that it is likely an upstream component of the *C. elegans* LCD pathway. To test this idea, we examined MYRF-1::GFP-AID animals also harboring a single-copy transgene expressing HSF-1 containing an R145A gain-of-function mutation, which promotes linker cell death even when upstream components are silenced (*8*). Indeed, we found that this mutation significantly reduces aberrant linker cell survival in MYRF-1::GFP-AID animals treated with auxin from ∼60% to ∼36% (Fig. 4A). Thus, MYRF-1 likely acts upstream of HSF-1.

**Fig. 4.**
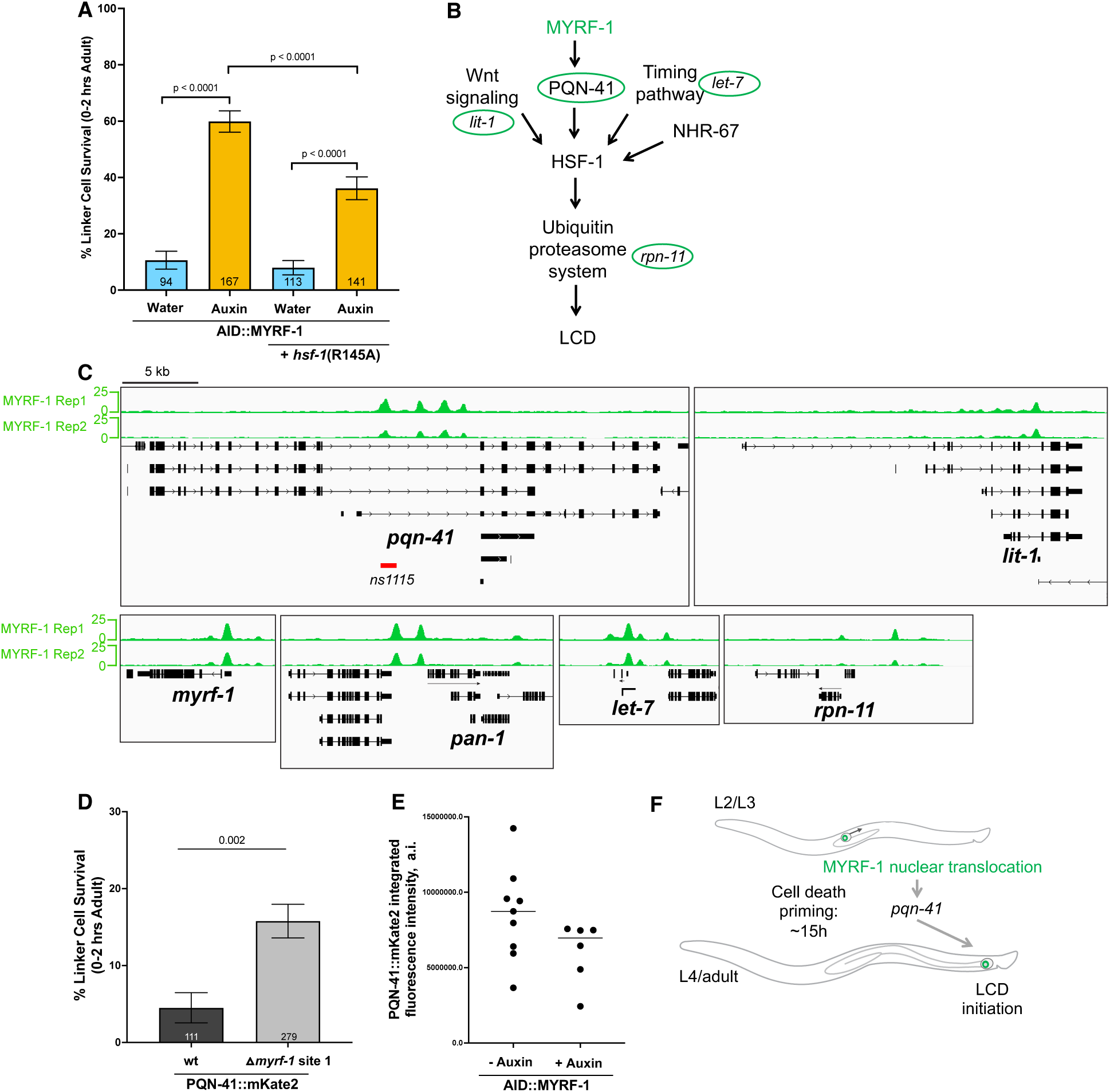
MYRF-1 controls expression of the polyQ encoding gene *pqn-41*. **(A-B)** HSF-1 gain-of-function suppresses MYRF-1-depletion LCD defect placing it upstream in the LCD genetic pathway. **(C)** ChIP-seq MYRF-1 binding peaks on LCD genes, highlighted in (B). **(D)** Deletion of MYRF-1 binding site 1 (*ns1115* allele) results in increased linker cell survival. **(E)** PQN-41::mKate2 levels are reduced upon MYRF-1 depletion. **(F)** Model of MYRF-1 priming for LCD.

To determine which LCD regulator may be the target of MYRF-1, we carried out a ChIP-seq study, using an N-terminally tagged GFP::MYRF-1, in early larvae in which MYRF-1 is localized to the nuclei of most expressing cells. Quality control data for this experiment are described in a parallel manuscript (Wu et al., in prep). Intriguingly, we found that MYRF-1 binds regulatory sequences of multiple LCD genes, including *pqn-41*, *lit-1*, *rpn-11*, *let-7*, *pan-1*, and *myrf-1* itself (Fig. 4B,C) (*1*, *8*, *10*).

To determine whether MYRF-1 binding regions are functionally relevant for linker cell death, we further characterized the *pqn-41* locus, encoding an endogenous polyglutamine protein. We identified four MYRF-1 binding peaks within a large *pqn-41* intron (Fig. 4C). Remarkably, CRISPR/Cas9 deletion of the most upstream binding site (*pqn-41(ns1115)*; Fig. 4C), results in aberrant linker cell survival (Fig. 4D). We also inserted the gene encoding mKate2 fluorescent protein just upstream of the *pqn-41* stop codon. The resulting fluorescent fusion protein is expressed in the linker cell, among other cells (fig. S4A), and auxin-induced MYRF-1 depletion reduces its expression in dying cells (Fig. 4E). Thus, MYRF-1 regulates *pqn-41* gene expression likely by direct binding to the *pqn-41* locus, and this binding is required for the fidelity of linker cell death.

The studies presented here reveal that the membrane-associated transcription factor MYRF-1 primes the *C. elegans* linker cell for death many hours and two developmental stages prior to cell death execution (Fig. 4F). In this *in vivo* setting, therefore, the commitment of the linker cell to die is temporally uncoupled from death onset. Mitochondrial priming was previously described in the context of apoptotic cell death, where it appears to determine the sensitivity of cultured cells to pro-apoptotic stimuli (*43*). The mechanism we describe here is, however, fundamentally distinct. First, priming occurs many hours prior to cell death onset, and not immediately before it occurs. Second, priming is regulated at the transcriptional level and not by changes to mitochondrial properties. Third, priming of linker cell death is developmentally programmed and occurs at high fidelity, unlike mitochondrial priming, which appears to be stochastic.

Our finding that MYRF-1 priming is mediated in part through *pqn-41*, a gene encoding a polyQ protein, is of particular interest, given the morphological similarities between LCD and polyglutamine-induced neurodegeneration in Huntington’s disease and related disorders (*3*). Of note, for both the linker cell and Huntington’s disease, it remains unexplained how broadly expressed factors affect the survival of highly specific cell types, an important mystery that may lie at the core of disease etiology. One possible solution is that MYRF-1 may activate different genes in different cells. Indeed, our finding that MYRF-2 is required for DD neuron remodeling but not for LCD suggests cell-specific targets.

Our studies also raise the possibility that human mutations in MYRF cause developmental abnormalities, in part, by aberrant control of cell death. Indeed, cell culling is an important aspect of developmental morphogenesis (*44*, *45*), and defects in cell death have been suggested to underly developmental errors in palate development and in other settings (*46*).

Why prime the linker cell for death? Our previous studies reveal that multiple upstream pathways control linker cell death onset. It is possible that priming allows sufficient time for the establishment of a signal integration mechanism that allows the linker cell to continually weigh the significance of multiple cues, initiating its own demise only at precisely the correct time. Initiation of linker cell death too early would lead to incomplete gonad elongation and sterility, whereas delaying linker cell death prevents gonad-cloaca fusion, blocking formation of an open reproductive channel. More broadly, our findings raise the possibility that irreversible developmental decisions may be encoded much earlier than their execution, a concept that has important implications for understanding disorders in which development goes awry.

## Supporting information

Data S1. All clusters markers

Data S2. Cluster15 Linker cell Average expression

Data S3. LCD and migration gene lists.

Data S4. Rare genes

## Acknowledgments

We thank Svetlana Mazel, Stanka Semova, Samer Shalaby and Songyan Han and the entire Rockefeller Flow Cytometry Resource Center for crucial assistant with cell sorting. We are grateful to the Rockefeller Genomics Resource Center, especially Connie Zhao, Helen Duan and Jackie Woodruff. We thank Matthew Paul and Thomas Carroll from the Rockefeller Bioinformatics Resource Center for organizing the single-cell sequencing analysis course. We would like to acknowledge the Electron Microscope Imaging Facility of CUNY Advanced Science Research Center for instrument use and scientific and technical assistance. We thank Maxime Kinet, Itai Toker, Piya Ghose, Katherine Stewart, Lena Kutscher, and Alex Edelstein and all members of the Shaham lab for discussions. We thank Betty Ortiz Bido for the PQN-41::mKate2 *C. elegans* strain. Some strains were provided by the CGC, which is funded by the NIH Office of Research Infrastructure Programs (P40OD010440). Claude Opus 4.1 was used for polishing of the first manuscript draft.

## Author contributions

Conceptualization: OY, SS

Data curation: OY, SL, PW, DFR, YL

Formal analysis: OY, SN, PW, YL

Funding acquisition: OY, CH, SE, SS

Investigation: OY, SL, YL, SN, TN, ST, SM, PW, DFR

Methodology: OY, SL, LBH, PW, DFR, SE

Project administration: OY, CH, SE, SS

Resources: LBH, DFR, SE, PW, CH

Software : OY, SL, PW

Supervision: SS, SE, CH

Validation: OY, SS

Visualization: OY, SL

Writing – original draft: OY, SS

Writing – review & editing: OY, SL, SS, LBH, YL, SN, TN, ST, SM, PW, DFR, CH, SE

## Data and materials availability

All data, code, and materials are available to the research community upon request. Raw, analyzed data and code will be publicly posted upon manuscript publication.

