## Supplementary material for "Non-apoptotic death of the *C. elegans* linker cell is primed by MYRF-1 activation of *pqn-41*/polyQ": Methods and Supplementary Figures

**MYRF-1 primes the migrating C. elegans linker cell for non-apoptotic cell death via a polyQ protein**

**The PDF file includes:**

Materials and Methods
Figs. S1 to S4

**Other Supplementary Materials for this manuscript include the following:**

Data S1 to S4

**Materials and Methods**

Strains

| Strain number | Genotype | Reference |
|---|---|---|
| OS15354 | *myrf-1(ns1102) II; qIs56 him-5(e1490) V, nsIs1045 X* | This study |
| OS15443 | *myrf-1(ns1102) II; pqn-41(syb7915) III; qIs56 him-5(e1490) V; nsIs1045 X* | This study |
| OS15501 | *pqn-41(ns1115) III; qIs56 him-5(e1490) V* | This study |
| OS15284 | *myrf-1(ybq133) II, qIs56 him-5 (e1490) V; nsIs1045 (X)* | Modified from (Qi) |
| OS15282 | *myrf-1(syb9904) II; qIs56 him-5 (e1490) V; nsIs1045 (X)* | This study |
| OS15316 | *myrf-1(syb9904) drSi28 II; qIs56 him-5 (e1490) V; nsIs1045 (X)* | This study |
| PXH10620 | *Flh-2(syb10620) III; him-5(e1490) V; nsIs650* | This study |
| OS15338 | *nsIS763 (II); him-5 (e1490) V; lam-2(qy20) X* | This study |
| OS15326 | *nsIS763 (II); epi-1(qy31) IV; him-5 (e1490) V* | This study |

Animal husbandry

*C. elegans* were maintained following standard protocols at 20°C on NGM plates supplemented with cholesterol and seeded with OP50 *E. coli* strain as food. For single cell experiments, animals were grown on high peptone NGM plates seeded with HB101 *E. coli* bacteria.

Preparation of synchronized larvae for single-cell dissociation

200,000 L4 larvae were used per dissociation reaction. On each experimental day, we processed 4-6 reactions totaling 800,000-1,200,000 larvae. Animals were expanded two weeks before the experimental day. Two strains were used: OS514 (*mig-24p::Venus; him-5(e1490)*) and OS8909 (*mig-24p::Venus; nhr-67p::mCherry; him-5(e1490)*). Control strains were Bristol N2 (unstained) for OS514, and OS514 (green only) and OS12667 (red only) for OS8909. At least two 15-cm high-peptone NGM plates containing HB101 *E. coli* and gravid hermaphrodites were bleached to produce sufficient L1 larvae for each experimental day. A standard hypochlorite bleaching protocol was used, with one 15-cm plate bleached per reaction. Embryos were incubated overnight at 20°C on a rotator and plated the next day at a density of 100,000 L1s per 15-cm plate. Two days later, L4 larvae were examined for linker cell position using a fluorescent dissecting scope and collected at two timepoints: late migrating (mid-L4) and dying (late-L4, at the tail).

Single-cell dissociation, FACS, and single-cell RNA sequencing

L4 larvae were collected by washing plates with M9 buffer. Two 15-cm plates (~200,000 animals, including ~67,000 males) were collected into a single 15-ml Falcon tube and washed ten times with M9 to remove all bacteria. The volume was reduced to 0.25 ml and animals were transferred to a 1.5-ml Eppendorf tube. Then, 0.5 ml of freshly thawed 2× SDS-DTT solution was added to each worm pellet. L4 larvae were incubated for no longer than 4.5-5 minutes (including 1-minute centrifugation and up to 1 minute of handling and pipetting)—this is a critical step. Worm pellets were washed five times with 1 ml ice-cold Egg buffer. Freshly dissolved pronase solution (15 mg/ml in Egg buffer; Sigma P8811-1G) was added, and samples were incubated on a rotator at 20°C for 5 minutes.

Larvae were mechanically disrupted using one of two methods: glass homogenizer or needle shearing. For glass homogenization, worms were transferred to a 2-ml glass homogenizer and vigorously homogenized for 4 minutes at room temperature using a type B pestle, followed by 6 minutes on ice. For needle shearing, samples were passed 30 times through a 25G×1" needle with a 3-ml syringe at 4°C. The supernatant was retained, and the pellet was processed with additional pronase incubation and needle extraction. Pronase treatment was quenched with ice-cold Egg buffer containing 0.1% BSA. Cells were washed three times by centrifugation at 5,000×g for 5 minutes at 4°C.

A Millex-SV 5.0-µm filter was pre-wetted with 1 ml Egg buffer containing 0.1% BSA and used to filter the cell suspension, which was resuspended in 500 µl ice-cold Egg buffer with 0.1% BSA. Residual cells were recovered with 1 ml Egg buffer containing 0.1% BSA. Cell density was measured using a hemocytometer. One worm pellet yielded $>1\times10^6$ cells/ml.

Collection tubes for FACS were pre-coated overnight at 4°C with Egg buffer containing 5% BSA. FACS was performed using a BD FACSAria equipped with a 70-µm diameter nozzle. DAPI (10 ng/ml final concentration; Thermo Fisher Scientific, Cat. #62248) was added to exclude dead cells. Gates were set to isolate DAPI-negative cells expressing GFP alone or both GFP and mCherry, depending on the strain. Sorted cells were counted and immediately processed for single-cell capture and library preparation using 10x Genomics Chromium Next GEM Single Cell 3' Reagents Kits v3.1 and v4. Each sample was loaded into a separate channel. Libraries were sequenced on an Illumina NextSeq 2000 system with 100-bp single-end reads. Raw sequencing data will be available upon publication.

<u>Single-cell RNA-seq data processing, quality control and integration</u>

For the five collected samples, sequencing reads were aligned to the *C. elegans* reference genome (WBcel235) using 10x Genomics Cell Ranger (v7.1.0) with default settings, including intronic reads. The resulting feature-barcode matrices were processed using CellBender (v0.2.1) for ambient RNA removal. Filtered matrices were imported into Seurat (v5) for downstream analysis. Cells with fewer than 200 genes (nFeature_RNA ≤200) or with >20% mitochondrial RNA content were excluded. The mitochondrial RNA threshold was set higher than standard to preserve dying cells. Doublets were identified using Scrublet and removed from the dataset. Overall, 78.9% of cells were retained post-filtering. The five individual filtered datasets were merged and integrated using Harmony to remove batch effects. Integration preserved temporal relationships between cells while eliminating batch differences between samples that potentially resulted from the use of different strains and sequencing kits.

<u>Cell type annotation</u>

Clustering was performed using UMAP. Clusters were annotated based on expression of established cell type-specific markers:

| **Cell Type** | **Markers** | **Key Citation (PMID)** |
|---|---|---|

| | | |
|---|---|---|
| **Neurons** | unc-119, rab-3, snb-1, unc-104 | PMID: 8582641, 9412487 |
| **Muscle** | myo-3, unc-54, pat-10 | PMID: 2583110, 6352051, 8106547 |
| **Germline** | pgl-1, glh-1, pie-1 | PMID: 9741628, 8943022, 1623519 |
| **Hypodermis** | lon-2, sem-2, cog-1, col genes | PMID: 9847238, 12482710, 10637627 |
| **Seam cells** | lin-11, sto-1 | PMID: 1970421, 18430929 |
| **Intestine** | gst-22, cld-9 | PMID: 20603539, 12819242 |
| **Pharynx** | myo-2, pha-4, phat-6/7 | PMID: 8244003, 7607089, 15238517 |
| **Coelomocytes** | unc-122 | PMID: 10545240 |

The linker cell cluster was identified based on expression of *hlh-3*, male specific genes *mab-3* and *dmd-3*, and known LCD genes *lin-29, let-7* and others (Fig. 1, S1C) (Mason et al., 2008; Rougvie & Ambros, 1995). LC identity was confirmed additionally confirmed by identity of cells isolated using both *mig-24p::Venus* and *nhr-67p::mCherry* reporters (fig. S1A,B). The linker cell cluster was subsetted without re-processing. Two outlier cells were removed, yielding a final linker cell cluster of 155 cells. As expected, we also annotated distal tip cells (DTCs) to cluster 1 and 5 (fig. S1D). Additionally, several other cell types were identified, including sperm, germ cells, and neurons, suggesting that the *mig-24p::Venus* reporter may be expressed at low levels more broadly than previously thought.

Pseudotime, differential gene expression and gene module analyses

Trajectory inference and pseudotime analysis were performed using monocle3 v1.3.4 (*23*) . We identified changing genes based on the pseudotime trajectory. Two subclusters were identified within the linker cell cluster. Their identify (migrating vs dying cells) was assigned based on the expression of markers (Fig. 2). Differential gene expression (DEG) analysis was performed by upending Monocle3 subclusers for cluster 15 (linker cell cluster) to Seurat and using Seurat's FindAllMarkers() function. Genes expressed in less than 10% of the total subcluster cells were excluded. Gene module analysis identified two main transcriptional modules. Genes identified by these three analyses were integrated to produce final Migration and LCD gene lists (Data S1).

GO enrichment analysis

Migration and LCD gene lists were loaded into GeneOntology.org (powered by Panther) and analyzed for biological function and cellular components. Significantly enriched categories were transferred to excel and R was used to make Dotplots (Fig. 2A)

Rare genes analysis

To identify genes with sparse expression patterns that show temporal specificity along the linker cell death trajectory, we developed a complementary analysis targeting genes expressed in 5-20% of cells. This was done using Claude Sonnet 4.5. Standard differential expression analyses are optimized for genes with higher expression frequencies and may miss biologically relevant genes with restricted temporal expression windows. Indeed, one of such genes, *viln-1* is expressed in just 8.4% of cells in cluster 15 (linker cell cluster) but is functionally important for LCD as shown by RNAi experiments (Fig. 2C).

We extracted normalized gene expression data from the linker cell Seurat object and identified "rare genes" as those expressed in 5-20% of cells with mean normalized expression >0.5 in expressing cells, ensuring detection of genes with meaningful signal rather than technical noise. For each rare gene, we performed a permutation-based test to assess whether expressing cells showed bias toward early or late pseudotime positions. Specifically, we calculated the mean pseudotime of cells expressing each gene and compared this to the distribution of mean pseudotime values obtained from 1,000 random samples of equal size. Genes were classified as early-biased if their mean pseudotime was less than the global median, or late-biased if greater than the median. P-values were calculated as the proportion of permutations showing equal or greater deviation from the global mean, and adjusted for multiple testing using the Benjamini-Hochberg method. Genes with adjusted p-value < 0.05 were considered to show significant temporal bias.

This approach identified 253 rare genes with significant pseudotime bias, including 79 early-biased and 174 late-biased genes, revealing temporal expression dynamics not captured by standard differential expression analysis. These rare temporally-restricted genes likely represent key regulators of specific stages in the linker cell death process.

<u>LC-specific RNAi</u>

RNAi was performed by feeding using available RNAi clones from the Ahringer library (*46*) . All clones were sequenced before use to confirm they targeted the correct gene. Bacterial cultures were grown overnight from single colonies in 3 ml LB with 100 µg/ml carbenicillin. Approximately 250 µl per plate was seeded onto dried RNAi plates. Seeded plates were incubated overnight at room temperature in the dark to induce RNAi expression. Feeding plates were prepared with 25 µg/ml carbenicillin and 1 mM IPTG (final concentrations). Plates were left in the dark for 4-7 days to dry.

We used a strain systemically deficient for RNAi (*rde-1(ne219)*) but carrying LC-specific *rde-1* rescue, rendering RNAi machinery functional specifically in linker cells. Additionally, the strain carried an LC reporter (*lag-2p::GFP*) and the *him-5* allele to increase male frequency. Gravid hermaphrodites were bleached on day 0, and eggs were incubated in M9 buffer overnight on a rotator. On day 1, synchronized L1s were plated onto seeded RNAi plates and scored two days later as 0-2 hour adults for the presence of surviving linker cells. At this timepoint, most LCs in wild-type animals have completed death. Empty vector was used as negative control, and RNAi targeting GFP served as positive control. All animals were scored for efficient GFP knockdown (≥60%) before experimental scoring.

<u>Linker cell-specific Auxin inducible degradation</u>

Exposure of *C. elegans* to the synthetic auxin analog K-NAA (1-napthaleneacetic Acid Potassium Salt), results in ubiquitination and subsequent proteasomal degradation of auxin inducible degron (AID)-tagged proteins in the presence of transgenically provided TIR1, the substrate recognition component of the E3 ubiquitin ligase complex (*40*, *47*) . Three MYRF-1 strains tagged with a degron sequence (*aid::gfp::myrf-1 [myrf-1(ns1102)]*; (*gfp::aid::myrf-1 [myrf-1(ybq133)],* (*mKate2::aid::myrf-1 [myrf-1(syb9904)])* were were crossed with a strain expressing TIR1 tagged with mRuby specifically in the linker cell (*nsIs1045 [mig-24p::TIR1-mRuby*] (see construction details below). K-NAA was dissolved in sterile water to prepare a 200 mM stock solution for 4 mM auxin treatment. OP50-seeded NGM plates were pre-coated with K-NAA to a final concentration of 4 mM. Dried plates were used immediately or next day, if used next days, pre-coated plates were kept in the dark. Synchronized populations of animals were transferred to K-NAA plates at specific stages. Control plates were pre-coated with sterile water.
For reversible auxin experiments, animals were washed off auxin plates with M9 buffer, spun down and transferred to water plates.

Linker cell survival scoring
Gravid hermaphrodites were bleached on day 0, and eggs were incubated in M9 buffer overnight on a rotator. On day 1, synchronized L1s were plated onto seeded plates. Two days later, molting males were picked into separate plates, and two hours later mounted onto 2% agar in M9 pads in 50 mM sodium azide. Surviving linker cells were found by fluorescent signal. Normal nuclear morphology was confirmed by DIC.

Fluorescence imaging
Fluorescence imaging was done using a Zeiss Imager M2 microscope equipped with an AxioCam MRm 1.4 camera and a 63x/1.4 NA oil immersion objective. For confocal images Zeiss LSM 900 inverted laser scanning confocal miscroscope with a 63x/1.4 NA oil objective was used. Animals were immobilized in 200 mM sodium azide on 2% agar in M9 pads.

Image quantification
GFP::MYRF-1 nuclear levels were scored using a confocal microscope and classified as high (bright nuclear signal), low (low nuclear signal), and none (no nuclear signal).

Generation of CRISPR/Cas9 alleles
*aid::gfp::myrf-1*
Ultramer oligo of 200nt carrying the original degron sequence flanked by 34bp myrf-1 homologies (IDT) was used as a single stranded donor to insert degron sequence right after the ATG and before the GFP sequence of the myrf-1 endogenous locus.
Ultramer sequence (homology arms, *degron*):
caaagaaccgatactttagcacacttcgaacATG*CCTAAAGATCCAGCCAAACCTCCGGCCAAGGCACAAGTTGTGGGATGGCCACCGGTGAGATCATACCGGAAGAACGTGATGGTTTCCTGCCAAAAATCAAGCGGTGGCCCGGAGGCGGCGGCGTTCGTGAAG*AGTAAAGGAGAAGAACTTTTCACTGGAGTTGTCC
crRNA: CACACTTCGAACATGAGTAA AGG(PAM)
The following mix was injected into young adult hermaphrodite gonads strain OS15321 (14 animals): tracrRNA (90 pmol), crRNA (95pmol), single stranded repair template (Ultramer), *rol-*

*6* plasmid (for F1 selection), purified Cas9 (30pmol). Animals were singled out and three days later F1 hermaphrodites that exhibit roller phenotype or are from plates with many roller animals were picked onto separate plates. Their progeny were genotyped using the following primers:
<u>WT allele – 164bp:</u>
Myrf-1_F: GCTTACAGAAGCCCTCCCAC
Myrf-1_int1_R: TGAGGTTGCAAAAATGTTCAGGT
Degron allele – 172bp
Myrf-1_F: GCTTACAGAAGCCCTCCCAC
Degron_R: CCACCGCTTGATTTTTGGCA
Degron insertion was confirmed by Sanger sequencing (Azenta).

*Deletion of MYRF-1 binding site within pqn-41 intron*
This site was deleted using two gRNAs and a repair oligo (ordered as Ultramer from IDT).
Upstream gRNA: AAAAGTCAATTGACGTAGTT CGG
Downstream gRNA: TGAGGAATACCCACTAAAAT TGG
Repair oligo:
AAAAGCTTCAAATTGAATTTAATAGGTTTCCGAACAATTGGAGTTTTCATGTGAAAA AACTTCAAATTTC

Injection mix containing both gRNAs, repair oligo, tracrRNA, Cas9 protein and coelomocyte::RFP selection marker was injected into 17 animals (strain OS15496). Because transgenic animals couldn't be detected, we allowed the plates from the injected animals to starve, then collected ~100 animals for genotyping using drop washing with lysis buffer.
Modified allele was detected with the following primers:
Mutant allele:
Pqn-41_myrf_site1_F AGTGGCCTGCACATTGTTCT
Pqn-41_myrf_site1_R CGAGAGAGGTCTGATGATTTCGT
WT product: 1133bp
Deleted allele product: 142bp

WT allele:
Pqn-41_myrf_site1_F AGTGGCCTGCACATTGTTCT
Pqn-41_myrf_site1_WT_R GTGAGGGAAAGAGCAGAGGG
Product: 229bp
Final sequence was confirmed by Sanger sequencing (Azenta).

<u>Construction of a linker cell-specific TIR-1 strain (nsIs1045[ mig-24p::TIR-1::mRuby]) for linker cell-specific AID</u>
mig-24p::TIR-1::mRuby (pLBH78) was made using Gibson assembly(*48*) . *TIR-1::mRuby* was amplified from pLZ31(*eft-3p::TIR-1::mRuby*) using primers homologous to the *mig-24* vector backbone (LBH124/125), and the *mig-24* promoter vector backbone was amplified from pLMK113 (*mig24p::mKate2*) using primers LBH15/16.
Assembled mig-24p::TIR-1::mRuby was injected at 20 ng/uL, with unc-122p::mcherry (40 ng/uL) as a coinjection marker, into the hermaphrodite gonad of qIs56[lag-2p::GFP] him-5(e1490) animals to generate transgenic lines as previously described (Mello, C. C. et al 1991).
The integrated array, *nsIs1045[mig-24p::TIR-1::mRuby (20ng/uL) unc-*

*122p::mCherry(40ng/uL)]*, was generated by exposing *qIs56 him-5(e1490); nsEx7419[mig-24p::TIR-1::mRuby (20ng/uL), unc-122p::mCherry (40 ng/uL)]* mutant animals to 33.4 mg/mL trioxsalen (Sigma T6137) and UV irradiation using a Stratagene Stratalinker UV 2400 Crosslinker (360 mJ/cm$^{2)}$ as previously described (*49*) . *nsIs1045* animals were outcrossed at least 4 times before being used in experiments.

Electron Microscopy

Animals were grown as described in the "Auxin-inducible degradation experiments" section. Two-hour adult males from water and auxin-treated AID::MYRF-1 plates were mounted on 2% agarose/M9 pads and lightly anesthetized with 25 mM sodium azide, then observed and imaged using fluorescence microscopy. Animals with surviving linker cells were recovered on food plates, then fixed within several minutes, stained, embedded in resin, and serially sectioned using standard methods (*50*) . Serial images were acquired using a Titan Themis 200 kV transmission electron microscope equipped with Cs image corrector. Image processing and analysis were performed using ImageJ and IMOD software.

ChIP-Seq of MYRF-1

Strain carrying MYRF-1, endogenously tagged with GFP at the N terminus, was used. Animals were synchronized and collected during late L1 stage when MYRF-1 is observed in the nucleus. Experimental details will be reported in the accompanying manuscript.

Statistical analysis

GraphPad Prism 10 (v10.6.0) and R (v4.3.2) were used for statistical analysis.

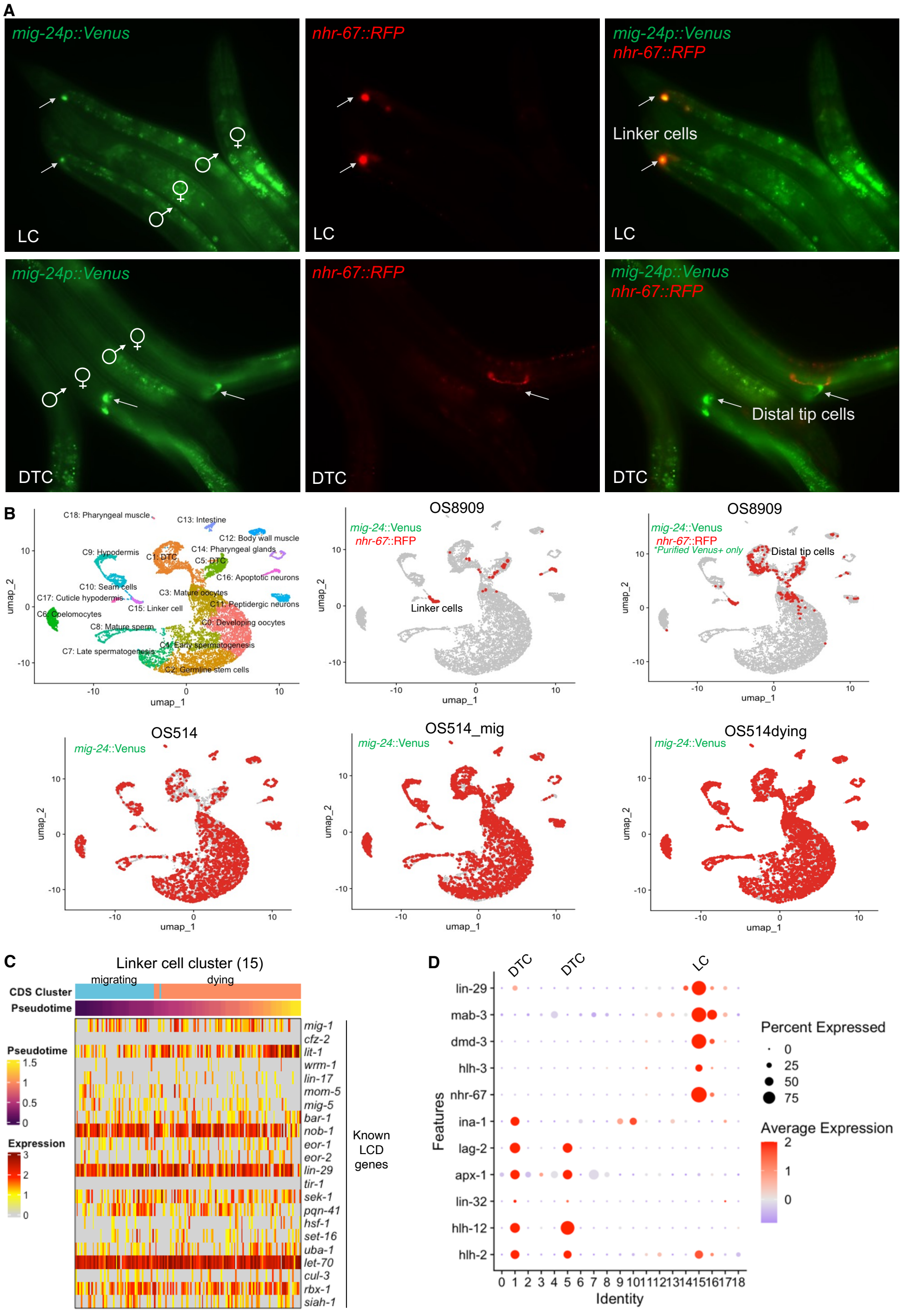


**Fig. S1. Identification of linker cell cluster.** **(A)** Demonstration of reporter specificity (LC – *mig-24p::Venus* + *nhr-67p::mCherry*; DTC – *mig-24p::Venus* only). **(B)** Identification of clusters and mapping of each dataset identity. **(C)** Linker cell cluster expresses all known LCD regulators. **(D)** Dotplot showing LC-specific markers in cluster 15, and DTC markers in clusters 1 and 5.

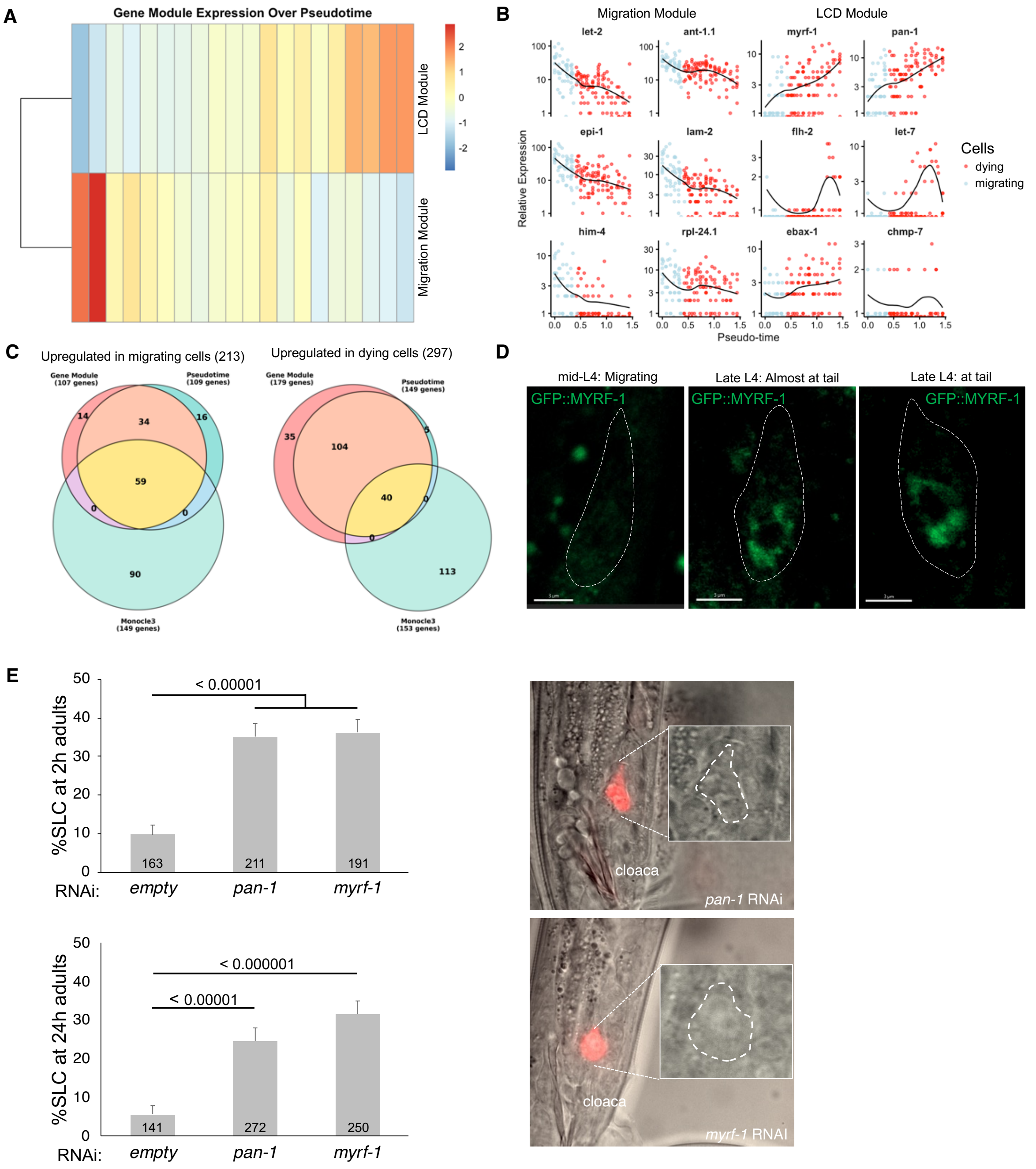


**Fig. S2. Additional single-cell RNA-seq analysis and validation of MYRF-1 and PAN-1 as novel LCD regulators. (A)** Two gene modules are detected in the linker cell cluster: LCD and Migration. **(B)** Examples of genes from these modules. **(C)** Venn diagram showing gene overlap from three different analysis to compile a final list of genes upregulated in migrating or dying cells. **(D)** GFP::MYRF-1 gets expressed and translocates to the nucleus when the linker cell arrives to the tail in the late L4 stage. **(E)** Linker cell abnormally survives upon myrf-1 and pan-1 RNAi knock-down in 2h and 24h adults. Representative images of 2h surviving cells.

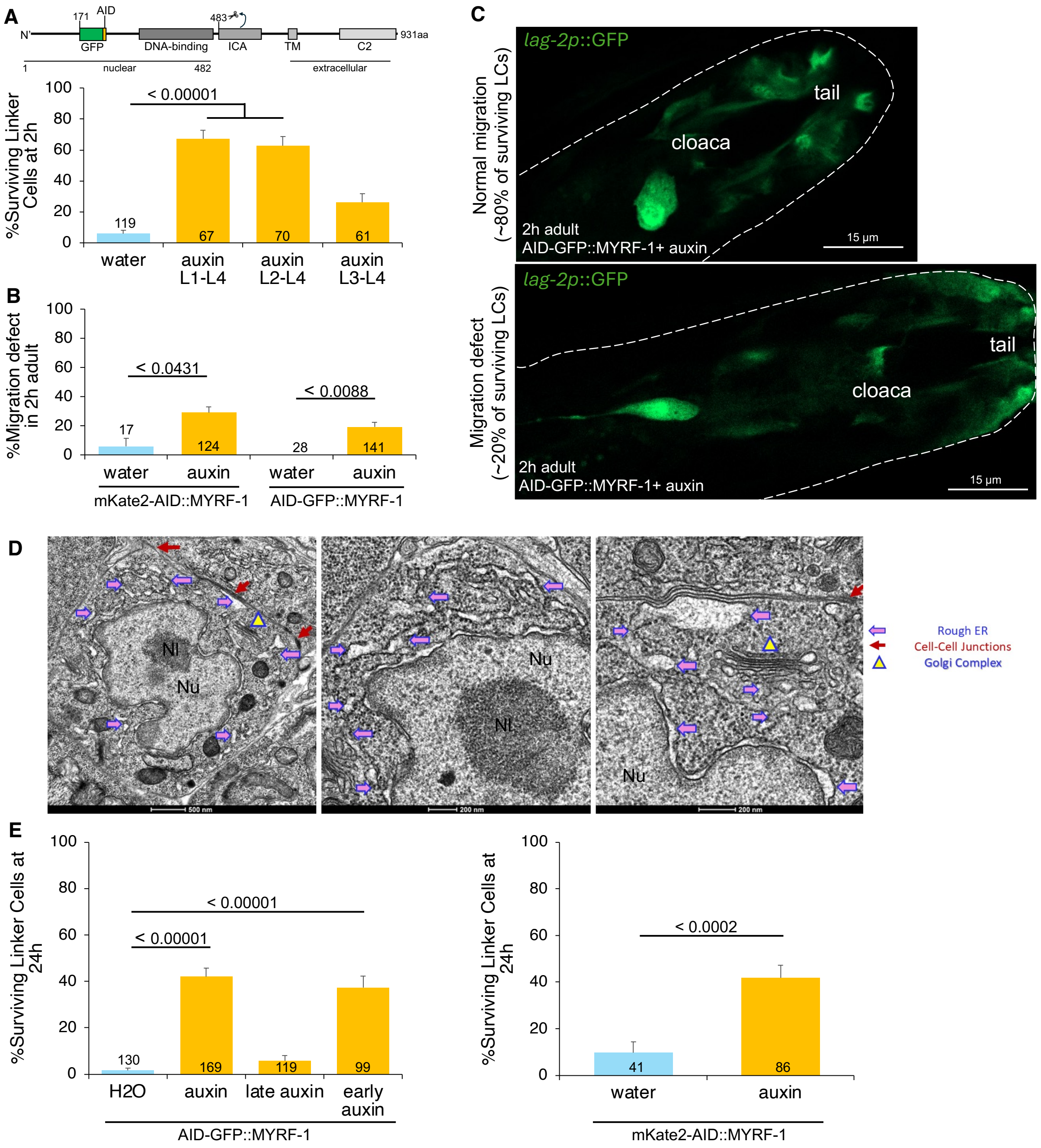


**Fig. S3. Linker cell migration is mostly unaffected upon MYRF-1 depletion, however its death is blocked in 2h and 24h adults. (A)** Auxin experiments using a third myrf-1 allele. **(B)** Quantification of migration defect in MYRF-1 depleted animals. **(C)** Representative confocal images of a normally migrating MYRF-1 depleted surviving linker cell and the one with a migration defect. **(D)** Electron micrographs of dying linker cells in control (water-treated) animals showing deformed nucleus (Nu) with nucleolus (Nl), ER swelling (purple arrows), normal Golgi (triangle) and cell-cell junctions (red arrows). **(E)** Linker cells survive 24h later in upon MYRF-1 depletion.

A

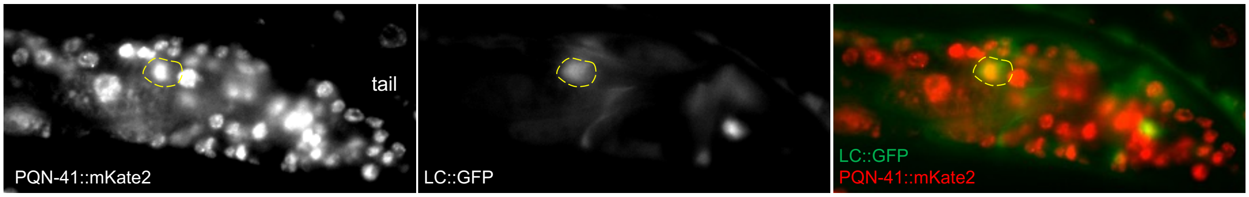


**Fig. S4. PQN-41 is expressed in the linker cell at late L4 stage.** *lag-2* promoter was used to drive GFP expression to label the linker cell.

**Data S1. (separate file)**
All clusters markers.

**Data S4. (separate file)**
Cluster15 Linker cell Average expression

**Data S3. (separate file)**
LCD and migration gene lists.

**Data S4. (separate file)**
Rare genes.
